# Gene Balance Predicts Transcriptional Responses Immediately Following Ploidy Change In *Arabidopsis thaliana*

**DOI:** 10.1101/795328

**Authors:** Barney Potter, Michael J. Song, Jeff J. Doyle, Jeremy E. Coate

**Author notes:** These authors contributed equally to this work (Barney Potter, Michael J. Song).

## Abstract

The Gene Balance Hypothesis postulates that there is selection on gene copy number (gene dosage) to preserve stoichiometric balance among interacting proteins. This presupposes that gene product abundance is governed by gene dosage, and that the way in which gene product abundance is governed by gene dosage is consistent for all genes in a dosage-sensitive network or complex. Gene dosage responses, however, have rarely been quantified and the available data suggest that they are highly variable. We sequenced the transcriptomes of two synthetic autopolyploid accessions of *Arabidopsis thaliana* and their diploid progenitors, as well as one natural tetraploid and its synthetic diploid produced via haploid induction, to estimate transcriptome size and gene dosage responses immediately following ploidy change. We demonstrate that overall transcriptome size does not exhibit a simple doubling in response to genome doubling, and that individual gene dosage responses are highly variable in all three accessions, indicating that expression is not strictly coupled with gene dosage. Nonetheless, putatively dosage-sensitive gene groups (GO terms, metabolic networks, gene families, and predicted interacting protein pairs) exhibit both smaller and more coordinated dosage responses than do putatively dosage-insensitive gene groups, suggesting that constraints on dosage balance operate immediately following whole genome duplication. This supports the hypothesis that duplicate gene retention patterns are shaped by selection to preserve dosage balance.

## Introduction

Gene duplication is prevalent in eukaryotic genomes, occurring with a frequency similar to that of single nucleotide substitutions (Lynch and Conery 2000, 2003; Tasdighian et al. 2017), and is a major contributor to genetic diversity and the evolution of novel traits (Lynch and Conery 2000) Most gene duplicates, however, are eventually pseudogenized and/or deleted from the genome, with an estimated half life for duplicated genes in plants of 17 MY (Lynch and Conery 2003). Following whole genome duplication (WGD, polyploidy) the majority of duplicated gene pairs (homoeologues) return to single copy (fractionate) in the process of diploidization (Langham et al. 2004; Schnable et al. 2011). A minority of duplicates from both small scale duplication (SSD) and WGD, however, escape this decay process, and are preserved over much longer periods of time. In *Arabidopsis*, for example, approximately 25 per cent of genes are retained in duplicate from the *α*-WGD ca. 32–43 MYA (Blanc and Wolfe 2004; Barker et al. 2009; Edger et al. 2018).

The retention or loss of redundant genes is not random. Certain classes of genes are preferentially retained in duplicated following WGD (Blanc and Wolfe 2004), and many of these same genes exhibit minimal duplication via SSD (e.g., tandem duplication, transposition) (Freeling 2009). This pattern, in which some classes of genes preferentially retain duplicates originating from WGD but retain few duplicates derived from SSD is referred to as “reciprocal retention” (Tasdighian et al. 2017).

Several models have been proposed to explain the long-term retention of duplicated genes (Panchy et al. 2016) including the evolution of new functions (neofunctionalization), division of ancestral function (subfunctionalization), selection on absolute dosage, and the Gene Balance Hypothesis (GBH) (Freeling 2009; Birchler and Veitia 2012; Papp et al. 2003). Among these, only the GBH provides an explanation for reciprocal retention. The GBH predicts that there is a fitness cost in disrupting the stoichiometric balance between proteins involved in coordinated networks (e.g., protein complexes and signaling cascades). By duplicating every gene in the network, WGD is thought to preserve this balance, and any subsequent gene losses would disrupt it. As a consequence, genes in these networks are retained together through the diploidization process via purifying selection to preserve balance. Conversely, duplicates arising from SSD disrupt balance in dosage-sensitive networks, and selection acts to purge them.

Three main lines of evidence support the GBH (Tasdighian et al. 2017; Freeling 2009; Hou et al. 2018; Edger and Pires 2009): 1) signaling cascades, regulatory networks and protein complexes that are known to be disrupted by unbalanced changes in protein abundance tend to exhibit reciprocal retention patterns; 2) reciprocally retained genes exhibit greater selective constraint on sequence evolution (lower Ka/Ks) and less divergence in expression patterns than non-reciprocally retained genes; and 3) reciprocally retained genes often exhibit deleterious phenotypes when over or under expressed—this latter piece of evidence often cited as the ultimate proof needed to demonstrate dosage sensitivity and confirm the GBH. However, demonstrating that a deleterious phenotype is induced by over- or under-expressing a gene provides evidence for dosage sensitivity at the protein level, but it does not necessarily follow that there exists dosage sensitivity at the level of gene copy number. Gene dosage differences alone do not produce the deleterious phenomena associated with imbalance; the genes must be transcribed and translated. If gene copy number is decoupled from the final protein concentration at the point of interaction (e.g., multi-subunit complex assembly), selection on preservation of gene copy number loses its power as an explanation for gene retention. Decoupling can occur through differential expression of genes encoding members of a dosage-sensitive complex, through differential stability of mRNAs encoding members of the complex, through differential translation of those mRNAs, or through differential stability of proteins.

Such decoupling is evident in response to polyploidy because not all genes show identical expression responses following duplication - whether measured at the level of transcript abundance (e.g. Pirrello et al. (2018); Hou et al. (2018); Guo et al. (1996); Riddle et al. (2006); Robinson et al. (2018); Stupar et al. (2007); Yu et al. (2010), additional references in Doyle and Coate (2019)) or protein abundance (Birchler and Newton 1981; Yao et al. 2011; Zhu et al. 2012; Soltis et al. 2016; Deng et al. 2017; Fan et al. 2017; Wang et al. 2017; Yan et al. 2017). Consequently, WGD does not necessarily preserve protein dosage balance, and the extent to which dosage responses following WGD are coordinated amongst genes encoding interacting proteins is unknown. To affect balance at the protein level, gene copy number minimally should be “felt” at the level of the transcriptome: For the GBH to have explanatory power as a force maintaining gene copy number, maintenance of transcriptomic balance is necessary, though not sufficient.

Therefore, the GBH predicts that: 1) genes in reciprocally retained networks exhibit changes in expression in response to WGD (they are not dosage compensated), and 2) that these changes are similar for all genes in the network (what we refer to as “coordinated responses”). Our previous study examined the relationship between duplication history and gene dosage responses at the level of transcription in *Glycine* neoallopolyploids (Coate et al. 2016). We showed that genes in reciprocally retained GO terms and metabolic networks showed more coordinated dosage responses than genes in non-reciprocally retained networks, consistent with gene dosage sensitivity. The Coate et al. (2016) study, however, was complicated by the fact that the observed expression patterns were the net result of WGD and hybridization, as well as by ca. 0.5 MY of post-WGD evolution. Additionally, we only measured relative expression levels (transcript concentrations) rather than absolute dosage responses. In fact, there remains very little data about the immediate dosage responses to “pure” doubling (autopolyploidy) (Spoelhof et al. 2017; Visger et al. 2019), and whether or not these dosage responses are consistent with the GBH.

Long-term patterns of gene retention and loss as predicted by the GBH rely on very simple assumptions that can be tested by synthetic polyploids, namely: gene duplication immediately alters gene expression and it does so in a coordinated fashion for genes encoding dosage-sensitive proteins. Synthetic polyploids allow us to see the instantaneous effects of gene duplication on gene expression, thereby testing this assumption that duplication alters expression. An additional expectation of the GBH is that that there should be low variation in transcript abundance among individuals for genes that encode proteins in dosage-sensitive complexes (Coate et al. 2016). This is because the stoichiometry of the complex would be disrupted when low-expressing alleles for some subunits are combined with high-expressing alleles for others. The current study, therefore, builds upon past work by using diploid/synthetic autotetraploid pairs of *Arabidopsis thaliana* (accessions C24 and Ws) and a tetraploid/synthetic diploid pair (Wa) to quantify transcriptome size, expression variance, and gene dosage responses in the first generations post-WGD in the absence of hybridization. We test whether there is an intrinsic, heritable difference between connected and non-connected genes and find that reciprocally retained gene groups immediately exhibit smaller and more coordinated dosage responses to changes in genome dosage (both WGD and genome halving) than their non-reciprocally retained counterparts.

## Materials and Methods

### Plant material

Gene dosage responses to ploidy change were quantified in two naturally occurring diploid *A. thaliana* accessions (C24 and Wassilewskija (Ws)) and colchicine-induced autotetraploids of the same accessions, as well as in one natural tetraploid accession (Warschau (Wa)) and a synthetic diploid generated by the Tailswap haploid induction system (Ravi and Chan 2010). All seeds were provided by Dr. Luca Comai. Seeds were sown on Sunshine #4 potting mix, cold stratified for four days, and grown in a growth chamber with 16/8 hour light/dark cycles at 21/18°*C*, respectively with ca. 125*µ mol*/*m*^2^/*s* light intensity.

### DNA/RNA Co-Extraction

Tissue was harvested from rosette leaves at the 10–12 leaf stage and DNA and RNA were coextracted using Qiagen AllPrep Universal kits. Extractions were performed on three to four individuals per accession. Nucleic acid yields were quantified by Qubit using DNA High Sensitivity and RNA Broad Range assays (Life Technologies). The size of the total RNA transcriptome (total RNA per unit of DNA) was estimated by the ratio of RNA to DNA.

### Flow cytometry

Endopolyploidy was quantified by flow cytometry. 50–75*mg* of leaf tissue was chopped with a razor blade in 600*µl* Aru buffer (Arumuganathan and Earle 1991). Suspended nuclei were filtered through a 20*µm* CellTrics filter (Partec), treated with RNAse (0.01*µg*/100*ml* of sample), and stained with propidium iodide (0.001*µg*/100*ml* of sample). Samples were analyzed on a FACSCanto II (BD Biosciences) flow cytometer to obtain counts per ploidy level and confirm the ploidy of the plants used for the study. Average ploidy level was determined by multiplying the fraction of events at a given ploidy level by the value of that ploidy level (i.e., 2, 4, 8, 16, 32, or 64), and summing the values for all ploidy levels.

### RNA-Seq

RNA-seq libraries were generated for each sample from 1–2*µg* of extracted RNA. To enable estimation of mRNA transcriptome size per unit of DNA, each sample was spiked with ERCC Mix 1 in proportion to the DNA/RNA ratio determined above, as described in Robinson et al. (2018). Libraries were generated using the Illumina TruSeq Stranded library prep kits. Libraries were multiplexed with 8–12 samples per lane and 100bp single end sequences were generated on an Illumina HiSeq 250 at the Cornell Biotechnology Resource Center’s genomics facility.

### RNA-seq data processing and analysis

Raw FASTQ files were trimmed and filtered to remove low-quality reads and technical sequences using Trimmomatic (Bolger et al. 2014) with the following settings: ILLUMINACLIP, TruSeq3-SE.fa:2:30, 10; LEADING, 3; TRAILING, 3; SLIDINGWINDOW, 4:15; MINLEN, 36. Filtered reads were aligned with HISAT2 (Pertea et al. 2016) to the *Arabidopsis* reference sequence (TAIR10) and to the ERCC reference. HTSeq (Anders et al. 2015) was used to determine read counts per gene. Fold changes in expression between ploidy levels and differentially expressed genes were identified using DESeq2 (Love et al. 2014). Fold-changes (FC; tetraploid/diploid) were calculated per transcriptome and per genome. Per transcriptome FC was calculated using the standard DESeq2 procedure which normalizes for *Arabidopsis* library size (total count of reads mapped to the *Arabidopsis* reference). To estimate FC per genome, *Arabidopsis* read counts were normalized by ERCC library size. ERCC-specific size factors were estimated by DESeq2 using the estimateSizeFactors function on ERCC read counts, and these size factors were then used to normalize DESeq2-based analysis of *Arabidopsis* read counts. FC per transcriptome is a measure of change in transcript concentration (what fraction of the total transcriptome is composed of transcripts from the gene in question). FC per genome is a measure of relative expression per gene copy or gene dosage response (change in expression per change in gene copy number).

Relative mRNA transcriptome size per genome (tetraploid/diploid) was estimated individually based on the FC estimates for each gene in the RNA-seq data set according to the equation:

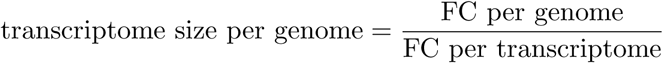

Reported values of transcriptome size per genome are the average of these individual estimates. Relative mRNA transcriptome size per cell was estimated by multiplying transcriptome size per genome by relative mean ploidy level (mean ploidy tetraploid/mean ploidy diploid)

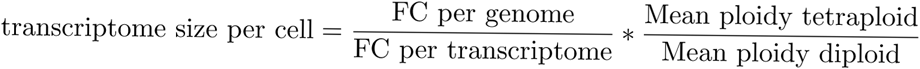

All scripts for data processing are available on GitHub.

## Results

### Classes of genes grouped by gene ontology and by metabolic network exhibit patterns of reciprocal retention

*Arabidopsis* genes were categorized as either singletons, WGD duplicates or SSD duplicates (including tandem, proximal or transposed duplicates) according to Wang et al. (2013). We then tested whether functionally related gene groups–gene ontologies (GO) or metabolic networks (Schlapfer et al. 2017)—exhibited patterns of reciprocal retention. As previously observed (Freeling 2009; Coate et al. 2016; Tasdighian et al. 2017), we found that both GO terms and metabolic networks with high retention following WGD tended to have lower retention of SSD (linear regression for GO terms, *slope* = −0.6972, *R*^2^ = 0.1839, *F* = 175.05, *df* = 1 and 777, *P <* 0.001; linear regression for metabolic networks, *slope* = 0.6667, *R*^2^ = 0.0886, *F* = 17.31, *df* = 1 and 178, *P <* 0.001; Fig. 1 a,b). This pattern is referred to as reciprocal retention (Tasdighian et al. 2017).

**Figure 1:**
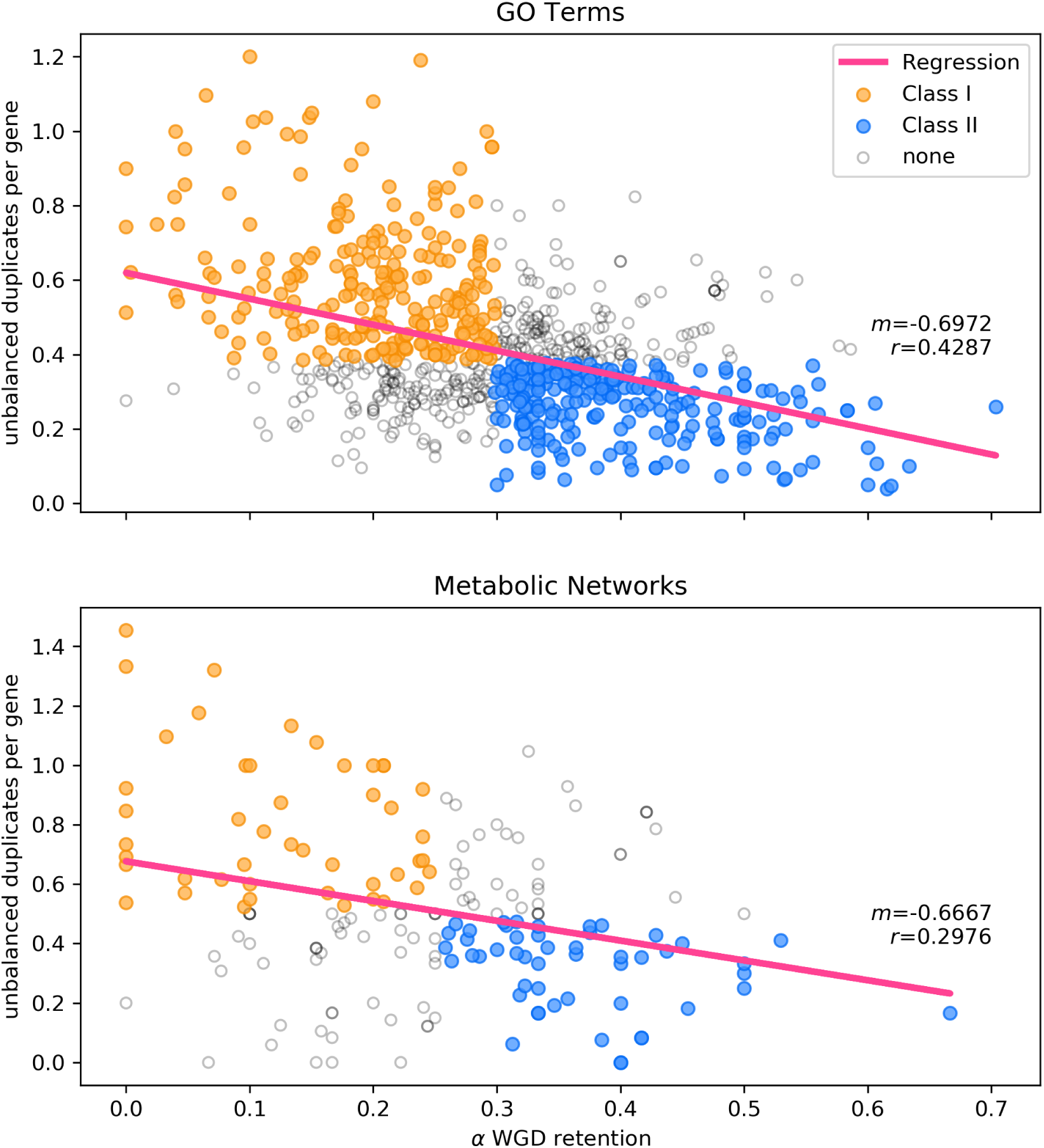
Reciprocal relationship between percentage of retained tandem duplicates and percentage of retained polyploid duplicate genes for GO classes (top) and metabolic networks (bottom).

To test whether the GBH explains these patterns of reciprocal retention, we grouped GO terms or networks into those that are putatively dosage insensitive (Class I; low WGD retention and high SSD retention, Fig. 1 yellow) and those that are putatively dosage sensitive (Class II; high WGD retention and low SSD retention, Fig. 1 blue) following the methods of Coate et al. (2016).

### Doubling the genome does not result in twice the total amount of transcripts per cell

The GBH depends on there being a strong correlation between gene dosage and transcript abundance (Coate et al. 2016). If gene dosage and transcript abundance are perfectly correlated for all genes then WGD would maintain a constant number of transcripts (transcriptome size) per genome resulting in a doubling of total transcripts per cell. We measured transcriptome size per genome and per cell to assess how closely transcript abundance correlates with gene copy number overall.

Both synthetic tetraploids (C24 and Ws) exhibited small but significant deviations in mRNA transcriptome size per genome relative to their diploid progenitors (*p <* 0.001; one-sample t-test; Table 1). Interestingly, the direction of change differed for the two accessions, with C24 exhibiting a small reduction in transcripts per genome (0.79-fold *±* 0.10 SD) and Ws exhibiting a small increase in transcripts per genome (1.19-fold *±* 0.06 SD). As with Ws, the natural tetraploid (Wa) exhibited slightly more transcripts per genome than its derived diploid (1.15-fold *±* 0.10 SD; *p <* 0.001; one-sample t-test). Thus, in none of the three accessions did genome doubling produce a simple doubling of transcripts, indicating that dosage responses per gene are variable, and deviate on average from a simple 1:1 dosage response.

**Table 1:** Summary statistics and Kruskal-Wallis tests for differences in PRV by Class for Gene Ontologies (GO), metabolic networks (AraCyc), Tasdigian et al. (2017) orthogroups (gene families), or Dong et al. (2019) structure based protein-protein interactions (S-PPI). N, number of functional groups included in the analysis.

| Grouping | Accession | N |  | Mean (SD) |  | Kruskal-Wallis |  |  |
| --- | --- | --- | --- | --- | --- | --- | --- | --- |
|  |  | Class I | Class II | Class I | Class II | X2 | df | p |
| GO | C24 | 188 | 199 | 0.494 (0.348) | 0.327 (0.253) | 35.59 | 1 | 2.44 x 10-09 |
|  | Ws | 185 | 191 | 0.348 (0.189) | 0.267 (0.133) | 26.341 | 1 | 2.86 x 10-07 |
|  | Wa | 186 | 194 | 0.233 (0.094) | 0.198 (0.089) | 16.952 | 1 | 3.83 x 10-05 |
| AraCyc | C24 | 29 | 41 | 0.428 (0.229) | 0.342 (0.223) | 3.3058 | 1 | 0.069 |
|  | Ws | 25 | 37 | 0.511 (0.567) | 0.262 (0.174) | 6.7835 | 1 | 0.0092 |
|  | Wa | 30 | 34 | 0.276 (0.164) | 0.181 (0.063) | 6.8824 | 1 | 0.0087 |
| Gene families | C24 | 141 | 652 | 0.407 (0.327) | 0.209 (0.211) | 62.531 | 1 | 2.62 x 10-15 |
|  | Ws | 127 | 618 | 0.334 (0.283) | 0.192 (0.187) | 39.95 | 1 | 2.60 x 10-10 |
|  | Wa | 149 | 650 | 0.356 (0.339) | 0.166 (0.188) | 54.2 | 1 | 1.81 x 10-13 |
| S-PPI | C24 | 7692 | 501 | 0.309 (0.318) | 0.223 (0.219) | 29.227 | 1 | 6.44 x 10-08 |
|  | Ws | 7416 | 484 | 0.236 (0.227) | 0.204 (0.193) | 9.0861 | 1 | 0.0026 |
|  | Wa | 8377 | 520 | 0.367 (0.466) | 0.242 (0.361) | 34.85 | 1 | 3.56 x 10-09 |

Notably, both synthetic tetraploids also exhibited reduced levels of endopolyploidy relative to their diploid progenitors (C24: *t* = *−*8.828, *df* = 4, *p <* 0.001; Ws: *t* = *−*3.416, *df* = 4, *p* = 0.027; two-sample t-test), such that mRNA transcriptome size per cell was, on average, significantly less than doubled in both accessions (*p <* 0.001; 1-sample t-test; Table 1). The size of the mRNA transcriptome per cell relative to the diploid progenitor was 1.14 *±* 0.14 for C24 and 1.49 *±* 0.08 for Ws. Thus, variable dosage responses and reduced endoreduplication interact to produce a smaller-than-expected transcriptome per cell on average, across all cell types and ploidy levels, although the effect in any given cell type is unknown.

The natural tetraploid, Wa, also exhibited a significantly lower level of endopolyploidy (*t* = *−*4.677, *df* = 7, *p* = 0.002; two-sample t-test) relative to its derived diploid, but the reduction was less extreme than in the derived tetraploids (average ploidy in the Wa tetraploid was 1.83-fold higher than in the diploid, compared to 1.37-fold higher in C24 and 1.25-fold higher in Ws). As a consequence, the derived diploid transcriptome per cell was roughly one half of the average natural tetraploid transcriptome (tetraploid:diploid: 2.11-fold *±* 0.18 SD).

### Individual gene dosage responses are highly variable, and many genes are dosage compensated

By quantifying transcriptome size we were able to estimate absolute dosage responses at individual loci (fold change in expression with a doubling of gene copy number). In all three accessions, dosage responses (change in transcripts per gene copy) were unimodally distributed around the estimate of overall transcriptome size, but with extreme values in each direction ranging from near silencing of expression with a doubling of gene copy number (a strong negative dosage effect) to a greater than 88-fold increase with a doubling in gene copy number (Fig. 2, Supplemental Fig. 1). 8.0%, 9.1% and 13.4% of genes deviated more than 2-fold from a 1:1 dosage response in Wa, Ws and C24, respectively.

**Figure 2:**
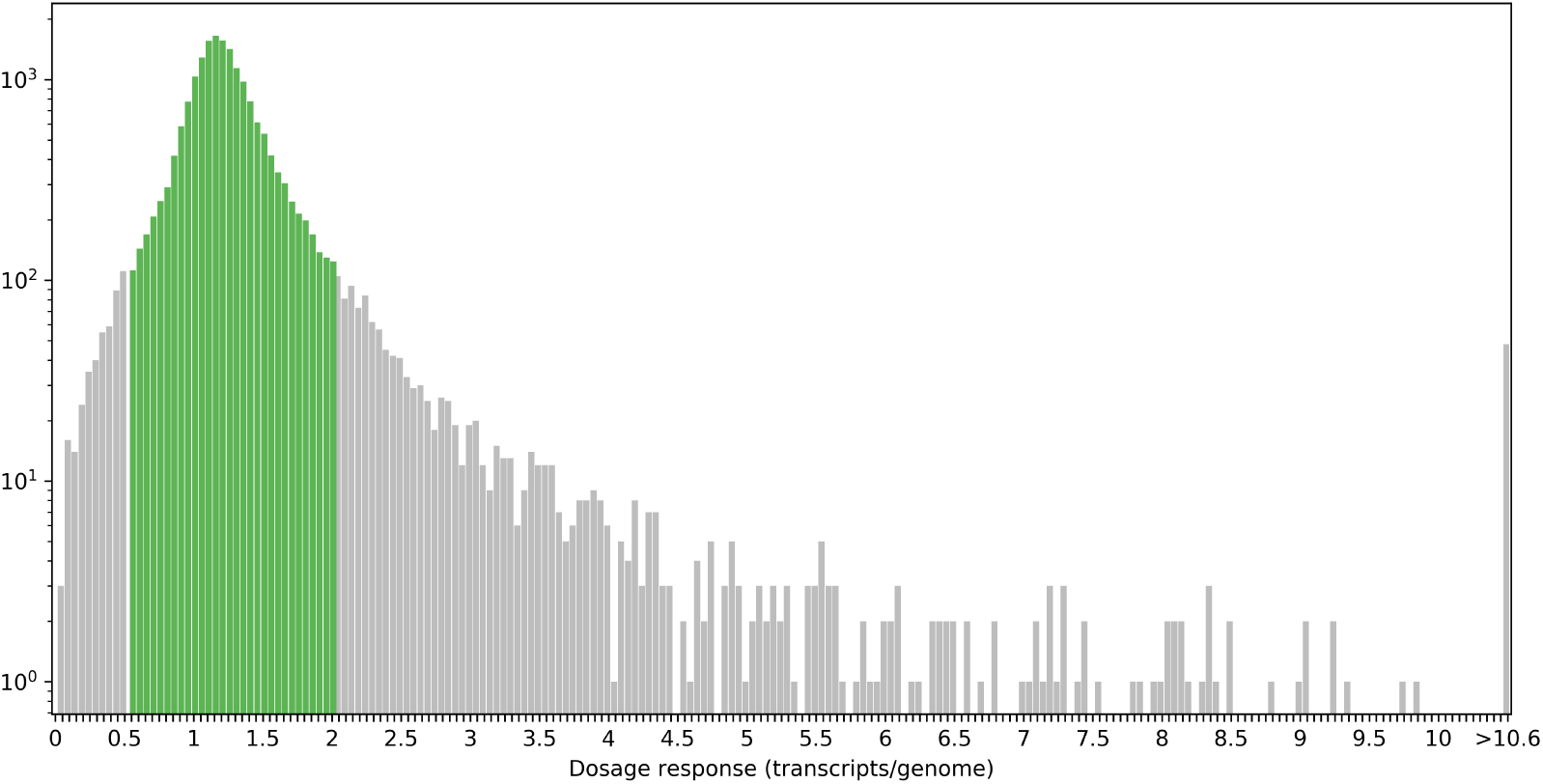
Distribution of gene dosage responses (transcripts per genome in the tetraploid divided by transcripts per genome in the diploid) in *Arabidopsis thaliana* accession Ws (N = 19,594 genes). A dosage response of 1 indicates a 1:1 dosage response (equal expression per gene copy or doubled expression per cell) in tetraploid vs. diploid. Dosage responses that differ by more than two-fold from a 1:1 dosage response are shown in grey (N = 1,789 genes; 9.1% of total). Values are cut off at 10 for display purposes, but 48 genes exhibit dosage responses *>*10 (maximum value = 88.7).

Additionally, many genes exhibit responses to WGD or genome halving consistent with dosage compensation (a halving of expression per gene copy, resulting in no change in expression per cell). For example, in Ws, the 95% confidence interval for transcripts per genome overlapped with 0.5 (dosage compensation) for 4,114 out of 19,594 genes for which we were able to estimate dosage responses (21%). 891 out of 21,259 genes (4.2%) and 7,061 out of 22,325 genes (31%) were dosage compensated in C24 and Wa, respectively. This is significant because dosage compensation decouples duplication from protein abundance. Thus, individual gene dosage responses are variable, and a large fraction of genes do not behave in a strictly dosage-dependent manner. Consequently, although the simplest way in which selection for maintaining balance among interacting proteins could drive reciprocal retention is if all genes exhibit 1:1 dosage responses (a 1:1 correspondence between transcript abundance and gene copy number, regardless of the mechanism of copy number change), this is not the case, regardless of whether the comparison is between synthetic polyploids and their natural diploid progenitors (C24 and Ws) or between a natural polyploid (Wa) and its synthetically derived diploid.

### Putatively dosage-sensitive gene classes exhibit coordinated dosage responses

Selection on dosage balance could still explain the reciprocal pattern of retention even given the lack of a uniform relationship between dosage and expression if all genes in a connected network have comparable, or coordinated, dosage responses (Coate et al. 2016). We tested if there are coordinated transcriptional responses to genome doubling for reciprocally-retained networks. Following the methods of Coate et al. (2016), for a given functional class (GO term) or metabolic network, we calculated the mean and coefficient of variation (CV; standard deviation divided by the mean) of dosage responses for all included genes. The CV, which we refer to as the Polyploid Response Variance (PRV) is a measure of the degree to which the dosage responses of genes within a network are correlated—a low PRV indicates strong coordination of dosage responses, whereas a large PRV indicates uncoordinated or variable dosage responses (Coate et al. 2016). We then looked to see if putatively dosage sensitive (Class II; reciprocally retained) networks or GO terms exhibit lower PRV than putatively insensitive (Class I; not reciprocally retained) networks or GO terms. Consistent with the GBH, PRV is lower for Class II than for Class I across all three polyploid-diploid pairs in all three categories (though the difference is not significant for metabolic networks in C24; (Table 1, Fig. 3).

**Figure 3:**
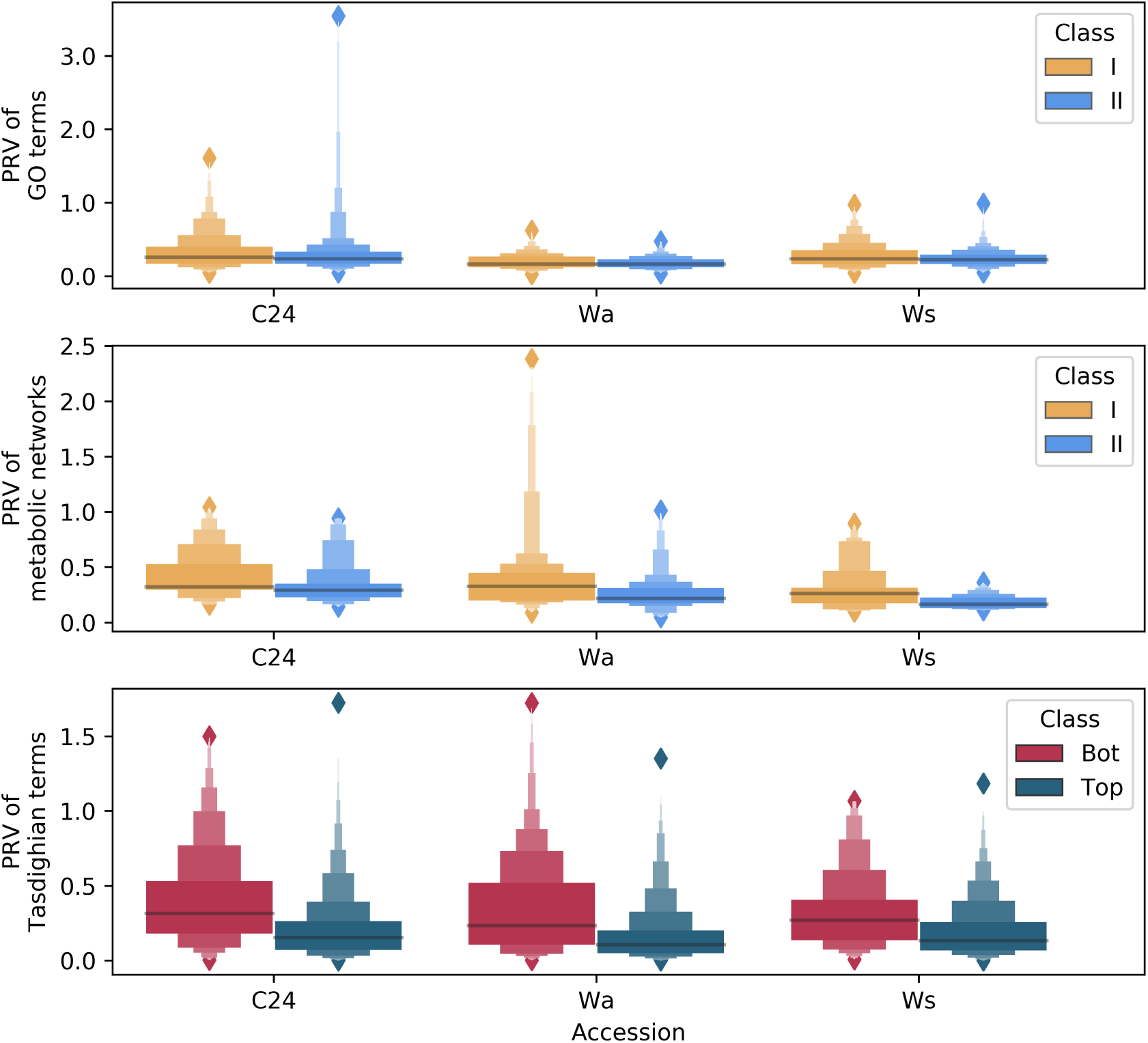
Polyploid response variance (PRV) by dosage sensitivity Class In C24, Wa and Ws for GO (top), metabolic networks (middle), and by reciprocal retention ranking of gene families (bottom; Tasdighian et al. 2017). Putatively dosage sensitive gene families (Class II) show lower average PRV than dosage insensitive gene families (Class I).

For the accessions with natural diploids and derived tetraploids (C24 and Ws), absolute dosage responses (fold-change in expression between tetraploids and diploids) were also significantly smaller on average in putatively dosage sensitive gene groups (Class II GO terms and metabolic networks) than in putatively insensitive groups (Class I GO terms and metabolic networks) (Fig. 4). In the natural tetraploid and derived diploid (Wa), dosage responses were also smaller for Class II functional groups, but the differences were not significant (Fig. 4).

**Figure 4:**
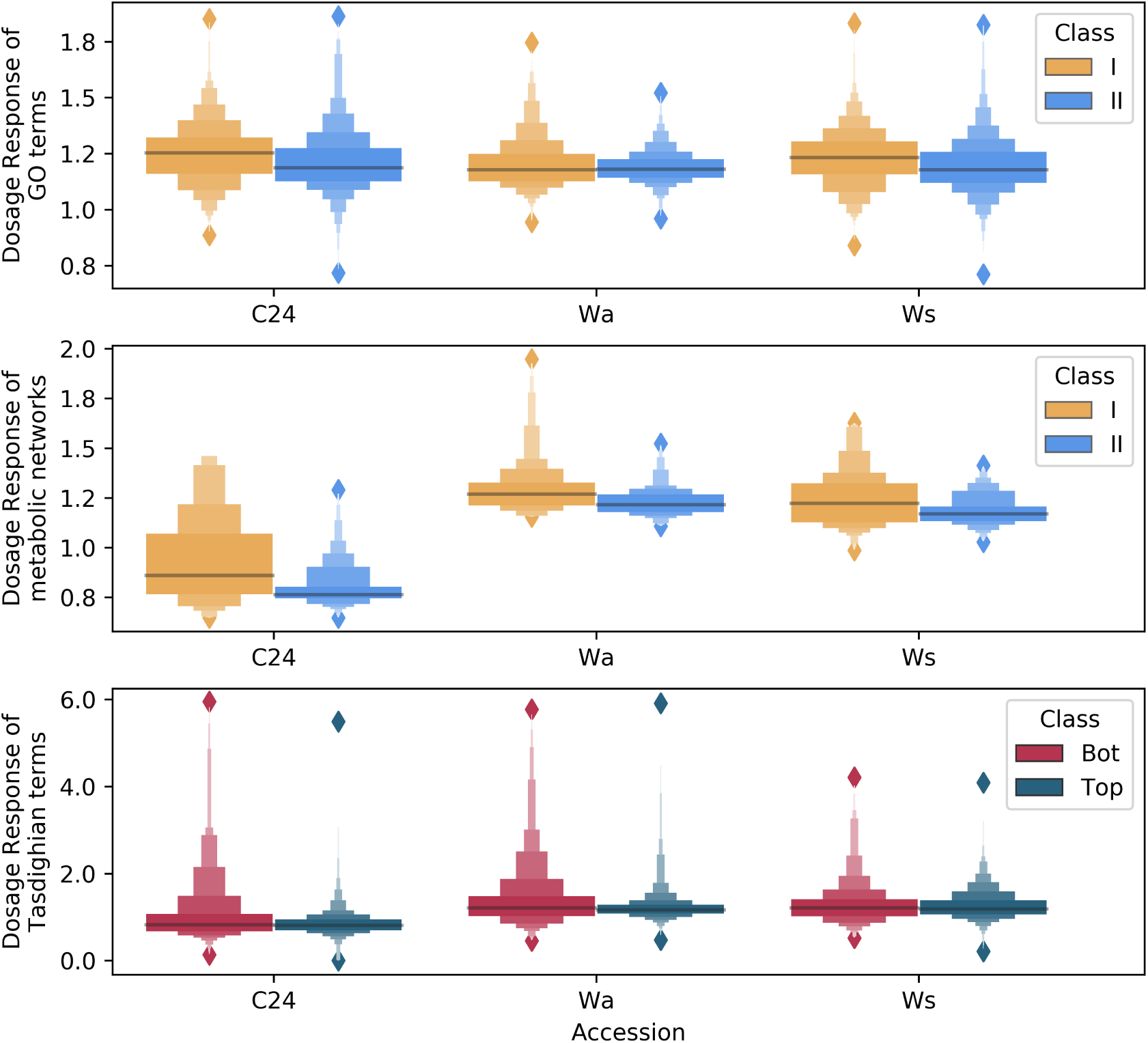
Dosage responses by dosage sensitivity Class In C24, Wa and Ws for GO (top), metabolic networks (middle) and by reciprocal retention ranking of gene families (bottom; Tasdighian et al. 2017). Putatively dosage sensitive gene families (Class II) show lower average dosage response than dosage insensitive gene families (Class I).

### Reciprocally retained gene families exhibit coordinated expression responses

Although there is a moderately strong pattern of reciprocal retention for GO terms, Tasdighian et al. (2017) have correctly pointed out that GO terms are sufficiently generic that many likely include both dosage-sensitive and dosage-insensitive genes. They argue that dosage sensitivity is better defined at the level of gene families as opposed to broad functional groupings. We therefore assessed if their 1000 most reciprocally retained gene families also exhibit lower PRV (more coordinated dosage responses) than do their 1000 least reciprocally retained gene families. We found coordinated expression responses consistent with the expectations of the GBH (Table 1, Fig. 3). Notably, the difference in PRV was more pronounced in this comparison than in the comparison of Class I vs Class II GO terms or metabolic networks, consistent with the Tasdighian et al. (2017) assertion that degree of dosage sensitivity is a characteristic of gene families and not necessarily a shared property of all genes of a broad functional category. In contrast to GO terms and metabolic networks, however, we did not observe smaller dosage responses in the top 1000 gene families than in the bottom 1000 gene families (Kruskal-Wallis tests: C24: *χ*^2^ = 2.95, *df* = 1, *p* = 0.086; Ws: *χ*^2^ = 0.01, *df* = 1, *p* = 0.903; Wa: *χ*^2^ = 2.65, *df* = 1, *p* = 0.103; Fig. 4).

### Dosage sensitive gene classes exhibit less variable expression levels across accessions

If dosage sensitive gene classes are under selection for coordinated expression of gene products, then these genes should exhibit similar expression levels across accessions within species to avoid expression imbalances resulting from recombining alleles (Coate et al. 2016). As expected, expression variance (EV) across accessions was smaller for Class II groupings (GO terms, metabolic networks and gene families) than for Class I groupings (Table 2, Fig. 5). In all groupings, this was true if we looked at EV among diploids, tetraploids, or diploids and tetraploids combined (Table 3).

**Table 2:** Summary statistics and Kruskal-Wallis tests for differences in dosage response by Class for Gene Ontologies (GO), metabolic networks (AraCyc), Tasdigian et al. (2017) orthogroups (gene families), or Dong et al. (2019) structure based protein-protein interactions (S-PPI). N, number of functional groups included in the analysis.

| Grouping | Accession | N |  | Mean (SD) |  | Kruskal-Wallis |  |  |
| --- | --- | --- | --- | --- | --- | --- | --- | --- |
|  |  | Class I | Class II | Class I | Class II | X2 | df | p |
| GO | C24 | 188 | 199 | 0.903 (0.240) | 0.845 (0.096) | 4.023 | 1 | 0.045 |
|  | Ws | 185 | 191 | 1.233 (0.117) | 1.206 (0.135) | 13.867 | 1 | 0.002 |
|  | Wa | 186 | 194 | 1.208 (0.101) | 1.194 (0.064) | 0.223 | 1 | 0.637 |
| AraCyc | C24 | 29 | 41 | 0.936 (0.236) | 0.799 (0.113) | 6.602 | 1 | 0.01 |
|  | Ws | 25 | 37 | 1.453 (0.582) | 1.230 (0.078) | 6.561 | 1 | 0.01 |
|  | Wa | 30 | 34 | 1.228 (0.136) | 1.177 (0.079) | 2.033 | 1 | 0.154 |
| Gene families | C24 | 141 | 652 | 1.162 (1.274) | 0.848 (0.326) | 2.946 | 1 | 0.086 |
|  | Ws | 127 | 618 | 1.735 (4.161) | 1.267 (0.504) | 0.015 | 1 | 0.903 |
|  | Wa | 149 | 650 | 1.880 (5.343) | 1.240 (0.529) | 2.653 | 1 | 0.103 |
| S-PPI | C24 | 7692 | 501 | 0.971 (1.264) | 0.822 (0.259) | 0.168 | 1 | 0.682 |
|  | Ws | 7416 | 484 | 1.346 (1.015) | 1.322 (0.425) | 3.72 | 1 | 0.054 |
|  | Wa | 8377 | 520 | 1.274 (1.045) | 1.300 (1.087) | 0.295 | 1 | 0.587 |

**Table 3:** Summary statistics and Kruskal-Wallis tests for differences in Expression Variance (EV) by Class for Gene Ontologies (GO), metabolic networks (AraCyc), Tasdigian et al. (2017) orthogroups (gene families), or Dong et al. (2019) structure based protein-protein interactions (S-PPI). N, number of functional groups included in the analysis.

| Functional group | Accessions | N |  | Mean ( SD) |  | Kruskal-Wallis |  |  |
| --- | --- | --- | --- | --- | --- | --- | --- | --- |
|  |  | Class I | Class II | Class I | Class II | X2 | df | p |
| GO | diploid | 174 | 190 | 0.274 (0.072) | 0.230 (0.055) | 33.396 | 1 | 7.52 x 10-09 |
|  | tetraploid | 174 | 190 | 0.304 (0.102) | 0.260 (0.062) | 16.007 | 1 | 6.31 x 10-05 |
|  | all | 174 | 190 | 0.291 (0.087) | 0.247 (0.056) | 23.605 | 1 | 1.18 x 10-06 |
| AraCyc | diploid | 26 | 37 | 0.292 (0.084) | 0.228 (0.060) | 9.01 | 1 | 0.0027 |
|  | tetraploid | 26 | 37 | 0.326 (0.124) | 0.251 (0.058) | 6.11 | 1 | 0.0135 |
|  | all | 26 | 37 | 0.312 (0.101) | 0.238 (0.056) | 8.43 | 1 | 0.0037 |
| Gene families | diploid | 77 | 501 | 0.327 (0.167) | 0.224 (0.123) | 30.16 | 1 | 3.98 x 10-8 |
|  | tetraploid | 77 | 501 | 0.356 (0.175) | 0.247 (0.133) | 31.495 | 1 | 2.00 x 10-8 |
|  | all | 77 | 501 | 0.344 (0.162) | 0.238 (0.110) | 34.276 | 1 | 4.78 x 10-9 |
| S-PPI | diploid | 5228 | 398 | 0.247 (0.141) | 0.202 (0.104) | 36.44 | 1 | 1.57E-09 |
|  | tetraploid | 5228 | 398 | 0.260 (0.169) | 0.252 (0.112) | 4.2141 | 1 | 0.04009 |
|  | all | 5228 | 398 | 0.253 (0.145) | 0.228 (0.090) | 2.1145 | 1 | 0.1459 |

**Figure 5:**
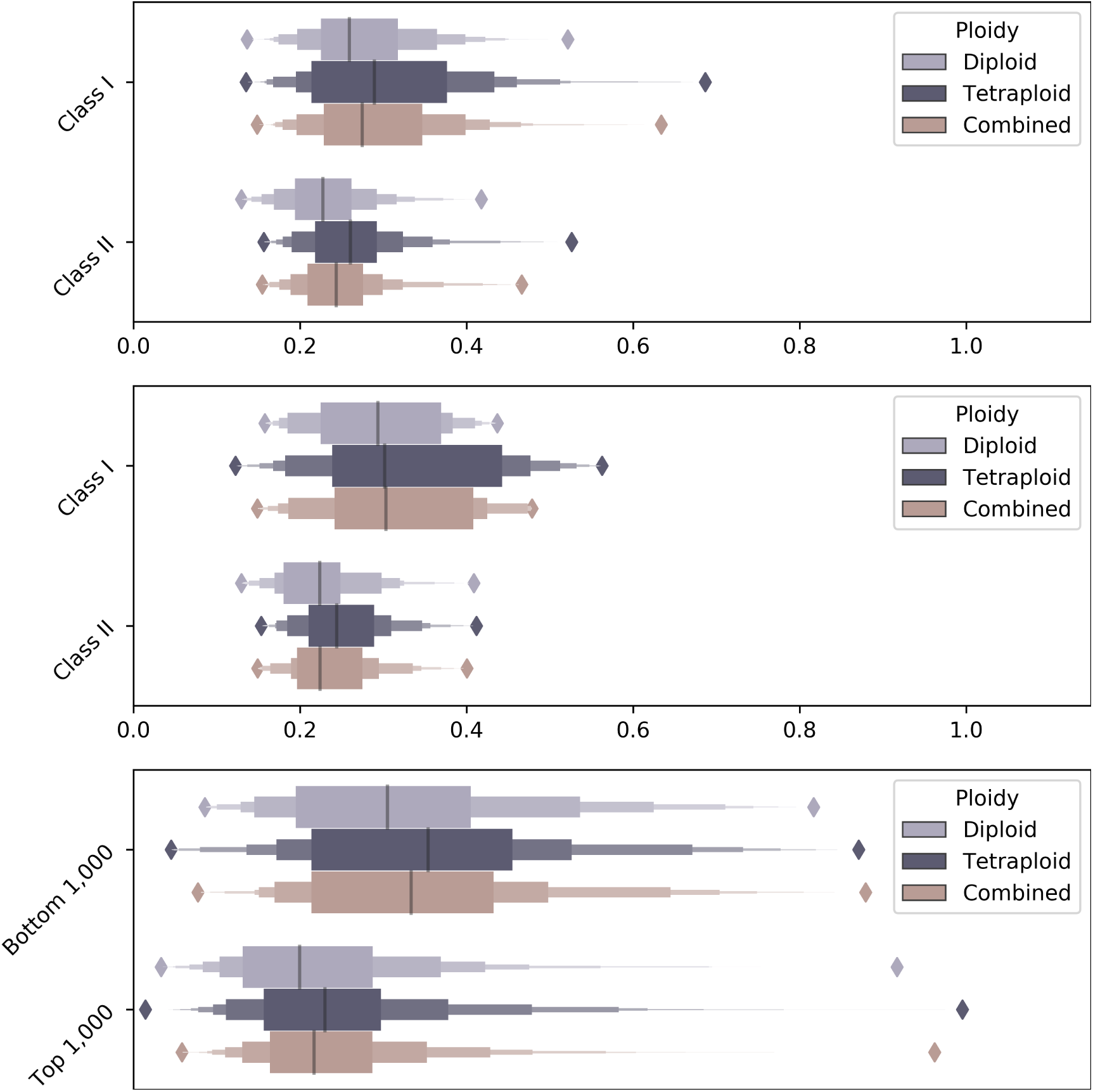
Expression variance (EV) by dosage sensitivity Class In diploids, tetraploids and diploids and tetraploids combined for GO (top), metabolic networks (middle) and by reciprocal retention ranking of gene families (bottom; Tasdighian et al. 2017). Putatively dosage sensitive gene families (Class II, Top 1000) show lower average dosage response than dosage insensitive gene families (Class I, Bottom 1000).

### Dosage sensitive predicted-interacting-protein pairs exhibit coordinated expression responses

Though Tasdighian et al. (2017) argue that dosage sensitivity is a property of gene families more so than of broader functional groups (e.g., GO terms), ultimately, dosage sensitivity presumably results from the need for stoichiometric balance between interacting proteins. In many cases, interacting proteins are members of the same gene family, but this is not always the case. We, therefore, next focused our analysis of expression patterns on protein-protein interactions. Using the top 1% ranked structure-based predicted protein-protein interactions (S-PPI) from Dong et al. (2019), we assessed whether the genes encoding interacting protein pairs exhibit a more coordinated expression pattern than random pairs of proteins. Surprisingly, on average, they do not. When separated by duplication history, however, we find that putatively dosage-sensitive protein pairs exhibit significantly lower PRV than do putative dosage-insensitive protein pairs (one or both encoding genes have lost their duplicate from the *α*-WGD and/or retain duplicates from SSD; Class I) (Table 1; Fig. 6). This reinforces the notion that not all protein-protein interactions are dosage sensitive, but that those protein-protein interactions that are dosage sensitive have evolved to maintain coordinated gene dosage responses. Looking at diploids and tetraploids separately, Class II protein-protein interactions also exhibit lower EV (Table 2).

**Figure 6:**
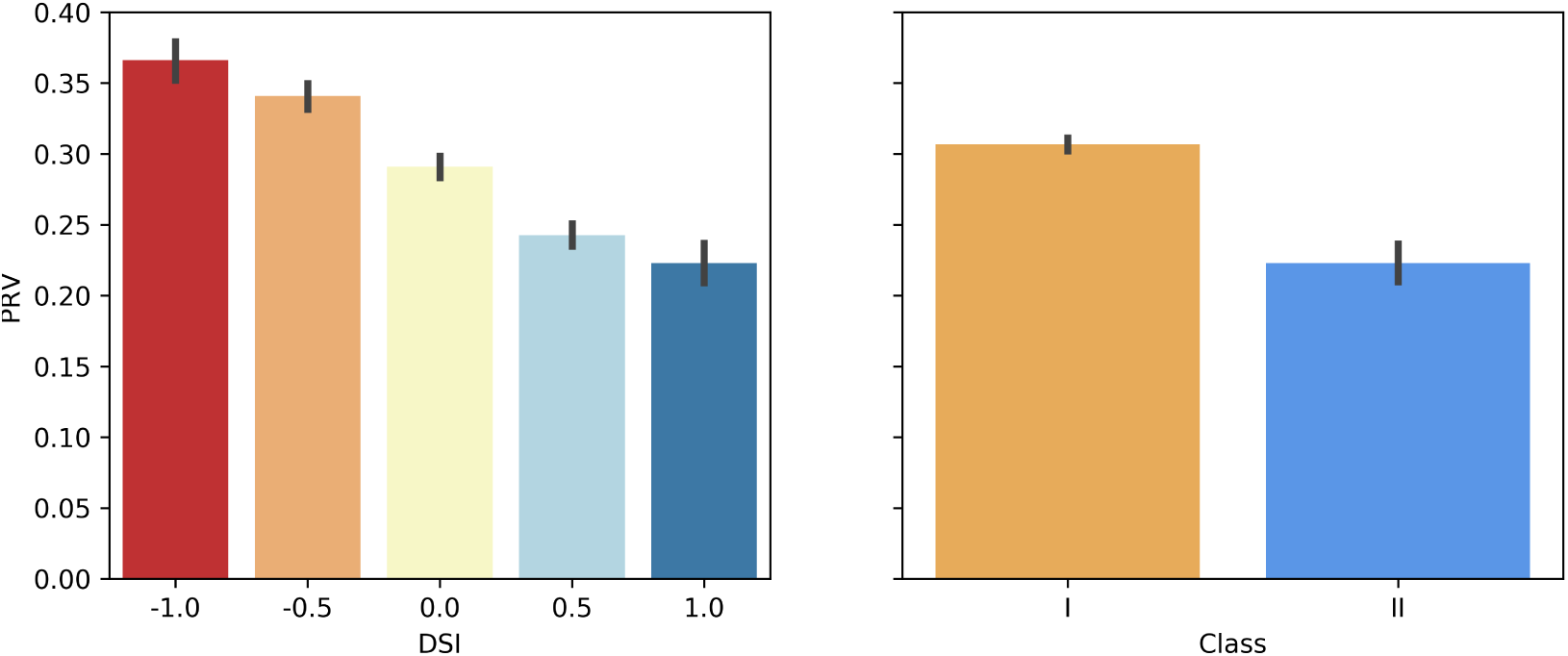
PRV by DSI *(left)* and Class *(right)* for predicted interacting pairs of proteins. Left, For each interacting protein pair, the duplication history of the encoding genes was used to calculated DSI, which is equal to WGD retention (1 if both genes have retained their *α* duplicate, 0.5 if 1 out of 2 has, 0 if neither has) minus small scale duplication (1 if both have been duplicated by small scale events, 0.5 if 1 out of 2 has, 0 if neither has). A DSI of 1 is evidence that the interaction is dosage sensitive, and decreasing values of DSI suggest decreasing levels of dosage sensitivity. Right, Class II is the same as DSI = 1 and Class I is everything else. PRV is calculated as described for GO terms and metabolic networks.

## Discussion

Although there is growing experimental support for selection on relative gene dosage (dosage balance) as a significant driver of the biased patterns of gene retention and loss following polyploidy, the logical steps between reciprocal retention and dosage sensitivity are just now being addressed (Tasdighian et al. 2017; Coate et al. 2016). Importantly, because the GBH assumes that selection operates to maintain relatively constant protein amounts among network members, it presupposes that gene dosage affects protein production. Examining the immediate transcriptional response to genome doubling, therefore, allows us to measure the extent to which expression level is driven by copy number and assess the potential for selection on gene dosage balance to shape the long-term evolutionary fate of genes.

We first estimated overall mRNA transcriptome size and found that it is not exactly doubled or halved with a doubling or halving of the genome, and that most genes do not exhibit simple 1:1 gene dosage responses. Hou et al. (2018) also observed slightly less than 1:1 increases in expression in a separate *Arabidopsis* ploidy series. Similar deviations from a simple 1:1 dosage response have been observed in leaf tissue of allotetraploid relatives of soybean (Coate and Doyle 2010), sepals of autotetraploid *Arabidopsis* (Robinson et al. 2018), and leaves of allotetraploid Tolmiea (Visger et al. 2019). Non-linear transcriptional responses to changes in gene dosage have also been observed following small scale duplications. For example, Konrad et al. (2018) observed greater than two-fold increases in expression following segmental duplication in *C. elegans*. In contrast, dosage compensation (minimal change in expression with gene doubling) has been observed in Drosophila yakuba, *D. melanogaster*, yeast and mammals (Qian et al. 2010; Rogers et al. 2017; Zhou et al. 2011). Zhou et al. (2011) for example, observed no differences in expression for 79% of 207 copy number variants in *D. melanogaster*.

Alleles share a common genomic address, and likely share more similar cis-regulatory environments than do paralogs (Gabaldón and Koonin 2013). Consequently, one might expect gene expression to be tightly correlated with allelic dosage. Yet even in the case of changes in allelic dosage, non-linear transcriptional responses are observed. For example, Springer et al. (2010) showed that 20% of allelic deletions did not result in a halving of protein abundance in yeast, with 3% of genes exhibiting dosage compensation. Thus, many genes deviate from a simple 1:1 relationship between gene dosage and transcript abundance, whether dosage is altered via allelic deletion/duplication, SSD or WGD. Furthermore, we observed different global transcriptional responses to WGD depending on accession. Similarly, Yu et al. (2010) found that *Arabidopsis* autotetraploids exhibited differences in transcriptional responses to WGD based on “ecotype” (genotype, accession), perhaps resulting from rapid cis-regulatory evolution and/or TE dynamics as observed, for example, in *Capsella* (Steige et al. 2015). Therefore, the simplistic assumption of the GBH—that WGD preserves protein dosage balance by equally increasing the abundance of all proteins—is not correct, and necessitates an assessment of whether or not stoichiometry is preserved by WGD for putatively dosage sensitive gene networks in the face of variable dosage responses.

Despite the observed disconnect between gene dosage and gene product amount, there might still be selection on gene dosage if genes in connected networks exhibit coordinated expression responses. Having estimated transcriptome size responses in both synthetic polyploid-natural diploid pairs and a synthetic diploid-natural polyploid pair, we were able to assay whether genes in reciprocally retained networks exhibit coordinated dosage responses. If dosage sensitivity explains patterns of retention long term, then there must be mechanisms to facilitate their co-regulation (Papp et al. 2003), and, by extension, coordinated responses to WGD.

Our data are consistent with this hypothesis. Reciprocally retained and, therefore, putatively dosage sensitive, gene groups (GO terms, metabolic networks, gene families, and predicted protein-protein interactions) exhibit less variable expression levels across accessions as well as more coordinated responses to changes in whole genome dosage. This pattern is consistent with our previous studies in *Glycine* (Coate et al. 2016), extending expression-level support for the GBH to autopolyploid systems. Thus, it appears that coordinated regulation within dosage sensitive networks is both independent of, and robust to, hybridization and the novel regulatory combinations that result.

A limitation of our previous study (Coate et al. 2016) is that it relied on natural tetraploids that are ca. 0.5 million years old. Thus, the expression patterns observed might reflect 0.5 million years (Bombarely et al. 2014) of independent evolution rather than (or in addition to) the immediate responses to genome doubling. The GBH, however, explains reciprocal retention as an “instant and neutral byproduct, a spandrel, of purifying selection” (Freeling 2009). For this to be true, coordinated expression responses need to be an instantaneous response to WGD. The comparison of induced polyploids to their isogenic diploid parents in the present study enabled us to assess if this is true, and demonstrates that reciprocally retained gene groups do, in fact, exhibit a higher degree of coordination in their dosage responses immediately following WGD.

It has been widely speculated that dosage constraints preserve duplicates in the short term, but that over longer evolutionary time periods, selection on gene dosage balance is relaxed, enabling the retained duplicates to subsequently subfunctionalize or neofunctionalize (Coate and Doyle 2011; Schnable et al. 2012; Conant et al. 2014; Coate et al. 2016; Gout and Lynch 2015). Under this scenario, one might expect to see more coordinated dosage-responses among reciprocally retained gene networks in nascent polyploids (where genes are under purifying selection to preserve dosage) than in older polyploids (where genes may be under relaxed selection on gene dosage with some having begun to diverge in function). Intriguingly, however, the degree to which dosage responses are more coordinated among Class II networks than among Class I networks is not discernibly more pronounced in the synthetic autotetraploids (current study) vs. natural allotetraploids (Coate et al. 2016). This could suggest that for most genes selection on gene dosage does not relax appreciably for more than a half-million years. This is consistent with observations that whole genome duplicates tend to diverge in expression more slowly than expected (Rodgers-Melnick et al. 2012; Tasdighian et al. 2017), and to diverge in expression and/or function more slowly than do small scale duplicates (Hakes et al. 2007; Qiao et al. 2018; Wang et al. 2011; Rodgers-Melnick et al. 2012; Defoort et al. 2019). If this is the case, performing equivalent analyses on older polyploids would help to resolve the timeline for when relaxation of selection on gene dosage occurs (e.g., cotton [*Gossypium hirsutum*], formed by allopolylpoidy 1–2 MYA).

Alternatively, or in addition, the lack of a stronger pattern in synthetic polyploids could be the result of deleterious (unbalanced) dosage responses arising at some loci in the nascent polyploids that are subsequently “corrected” by selection in polyploid lineages that survive the initial shift in genome dosage. We demonstrate that Class II gene groups show more coordinated dosage responses than do Class I groupings, but there is still considerable variation in dosage responses within Class II groups, some of which could represent unbalanced and, therefore, deleterious expression patterns that are rectified by purifying selection over subsequent generations.

Our study expanded the scope of Coate et al. (2016), which looked at GO and metabolic networks, by also assessing the top and bottom dosage sensitive gene families from Tasdighian et al. (2017), which the authors argue reveals a clearer pattern as dosage sensitivity is better measured at the level of gene families than broad functional groups where direct interactions between genes are less certain. Consistent with their assertion, we observed highly significant reductions in both PRV and EV in the top 1000 gene families relative to the bottom 1000 gene families (Figs. 3 & 5, Tables 1 & 3), and the differences were generally more pronounced than those observed between class II and class I GO terms or metabolic networks.

Likewise, with the recent publication of an *Arabidopsis* predicted protein-protein interaction network (Dong et al. 2019), we were also able to investigate the GBH on more explicitly interacting gene products as opposed to the indirect estimates provided by GO terms, metabolic networks or gene families for which the gene products do not necessarily interact. In all cases, we found a strong, consistent pattern of coordinated gene dosage responses across dosage sensitive groups, networks, and interacting protein pairs.

A prediction of the GBH is that genes in dosage-sensitive networks will be co-regulated, and Papp et al. (2003) provided evidence that this is in fact the case in yeast. We extend upon this observation to show that these genes are not only co-regulated within and across genomes at a given ploidy level, but that they are co-regulated in terms of their response to WGD.

One possible explanation for this surprising observation is that connected genes have evolved to share the same cis-regulatory element(s) (i.e., transcription factor binding sites), whereas unconnected genes have not. By sharing the same cis-regulatory modules, connected genes will be regulated by the same complement of transcription factors, which would facilitate co-regulation and, therefore, be favored by selection to preserve balance in dosage-sensitive complexes or signaling cascades. Sharing common cis-regulatory elements, therefore, would explain why such genes tend to be co-regulated as well as why they show coordinated dosage responses. Any change in the expression of these shared transcription factors in response to WGD (abundance increases, decreases, or stays the same) would affect all members of the connected network equally, enabling coordinated responses to WGD. Because Class I gene groups (GO terms, metabolic networks, etc.) are not dosage-sensitive, there is no selection favoring the acquisition of shared cis-elements. Consequently, they are more likely to be regulated by different sets of TFs, which themselves might exhibit different responses to WGD. As a result, Class I target genes (the unconnected genes) show less-coordinated expression responses to WGD. Consistent with this hypothesis, Taggart and Li (2018) demonstrated that proteins in complexes with obligate stoichiometry are produced in proportion to their dosage and concluded that their expression levels are hard-wired by cis-regulatory sequences.

A related explanation could be that dosage-sensitive gene groups reside in common chromatin contexts that coordinate expression. Though *Arabidopsis* generally lacks TADs ((Liu et al. 2017)), it does have various other chromatin interaction domains, including local chromatin loops (Liu et al. 2017), an intra- and inter-chromosomal structure termed the KNOT (Grob et al. 2014; Grob and Grossniklaus 2017), A and B compartments (Grob et al. 2014), “positive strips” and TAD-like structures (Wang et al. 2015), all of which correlate with specific expression profiles. Nuclear pore complexes are subnuclear compartments that are thought to be involved in organizing chromatin domains and thereby regulating transcription (Sun et al. 2019). Selection could favor the arrangement of genes from dosage-sensitive complexes into common chromatin domains, potentially mediated by nuclear pore complexes, to ensure co-regulation. Xie et al. (2019) showed that TADs and A/B compartments are largely conserved across related *Brassica* species. To the extent that these structures also persist after WGD events, these too could facilitate coordinated gene dosage responses. Notably, Xie et al. (2019) found that duplicates retained from the whole genome triplication event in *Brassica* were more likely to be colocalized in 3D chromatin domains. Thus, colocalization in chromatin domains is associated with both co-regulation and elevated duplicate retention following WGD. These observations are consistent with the notion that dosage-sensitive genes have evolved to be co-regulated via colocalization in shared chromatin domains, which in turn favors retention of balanced gene duplicates.

Transposable elements (TEs) can also provide an innate mechanism of expression coordination following polyploidization. Zhang et al. (2015) showed that WGD induces methylation in Class II TEs, which suppresses expression of nearby genes. They proposed that this could minimize deleterious gene dosage effects. Perhaps selection has favored the arrangement of dosage-sensitive gene networks in close proximity to DNA elements facilitating coordinated suppression of gene expression within dosage-sensitive networks post-WGD.

This TE-based mechanism would be consistent with our observation that putatively dosage sensitive GO terms and metabolic networks (but not gene families or interacting protein pairs) tend to show smaller average dosage responses (Fig. 4). It has been proposed that partial dosage compensation is due to selection to minimize disruption of balance by minimizing transcriptional change in response to change in gene dosage. Katju and Bergthorsson (2018) explain that this could be due to the relatively higher fitness cost of duplicating highly expressed genes and its associated increase in transcript abundance. Likewise Qian et al. (2010) describe expression reduction as a special class of subfunctionalization that could help explain the retention of duplicates.

These two studies provide a useful framework for why dosage sensitive genes have evolved to have smaller dosage responses (to minimize disruptions to balance from small scale duplications) and therefore as a corollary, smaller dosage responses offer further evidence that these genes are dosage sensitive. Qian et al. (2010) proposed that selection favors regulatory mutations that reduce expression. However, we observe smaller dosage responses for Class II genes in the first generations post-WGD, making it unlikely that post-duplication mutations are the cause. Epigenetic suppression resulting from the methylation of TEs could, therefore, be a plausible mechanism. It would be interesting to determine, therefore, if Class II genes are preferentially located in the vicinity of TEs.

Finally, while our study indicates that reciprocally-retained gene groups exhibit transcriptional responses consistent with the Gene Balance Hypothesis, it does not address whether these coordinated transcriptional responses produce coordination at the level of protein abundance. Multiple layers of post-transcriptional gene regulation could potentially result in imbalance at the protein level despite maintenance of balance at the gene dosage and/or transcriptional levels. Performing similar analyses to those presented here, but that incorporate ribosome profiling (Taggart and Li 2018) and/or quantitative proteomic data, would be necessary to fully assess whether protein dosage is sufficiently linked with gene dosage for selection to act on gene copy number to preserve balance in protein complexes and signaling cascades. Nonetheless, although quantifying proteins would provide the most direct evidence for this important assumption, any influence of gene dosage on protein abundance is presumably mediated by transcription, so the fact that the expected patterns are observed at the level of transcription attests to the efficacy of even these more indirect approaches and provides an important layer of support for the GBH.

## Acknowledgements

NSF grant 1257522 awarded to JEC and JJD. Reed College Summer Undergraduate Research Fellowship to BIP. XSEDE allocation TG-BIO170018 granted to JEC, MJS, and BP

## Author Contribution

JEC, JJD, and BIP designed the experiment. JEC and BIP performed the research. JEC, BIP, and MJS analyzed the data. JEC, BIP, MJS, and JJD wrote the manuscript.

## Literature Cited

Anders, S., P. T. Pyl, and W. Huber, 2015. Htseq-a python framework to work with high-throughput sequencing data. Bioinformatics 31:166–169.

Arumuganathan, K. and E. Earle, 1991. Nuclear dna content of some important plant species. Plant Molecular Biology Reporter 9:208–218.

Barker, M. S., H. Vogel, and M. E. Schranz, 2009. Paleopolyploidy in the brassicales: analyses of the cleome transcriptome elucidate the history of genome duplications in arabidopsis and other brassicales. Genome Biology and Evolution 1:391–399.

Birchler, J. A. and K. J. Newton, 1981. Modulation of protein levels in chromosomal dosage series of maize: the biochemical basis of aneuploid syndromes. Genetics 99:247–266.

Birchler, J. A. and R. A. Veitia, 2012. Gene balance hypothesis: connecting issues of dosage sensitivity across biological disciplines. Proceedings of the National Academy of Sciences 109:14746–14753.

Blanc, G. and K. H. Wolfe, 2004. Functional divergence of duplicated genes formed by polyploidy during arabidopsis evolution. The Plant Cell 16:1679–1691.

Bolger, A. M., M. Lohse, and B. Usadel, 2014. Trimmomatic: a flexible trimmer for illumina sequence data. Bioinformatics 30:2114–2120.

Bombarely, A., J. E. Coate, and J. J. Doyle, 2014. Mining transcriptomic data to study the origins and evolution of a plant allopolyploid complex. PeerJ 2:e391.

Coate, J. E. and J. J. Doyle, 2010. Quantifying whole transcriptome size, a prerequisite for understanding transcriptome evolution across species: an example from a plant allopolyploid. Genome Biology and Evolution 2:534–546.

Coate, J. E. and J. J. Doyle, 2011. Divergent evolutionary fates of major photosynthetic gene networks following gene and whole genome duplications. Plant Signaling & Behavior 6:594–597.

Coate, J. E., M. J. Song, A. Bombarely, and J. J. Doyle, 2016. Expression-level support for gene dosage sensitivity in three glycine subgenus glycine polyploids and their diploid progenitors. New Phytologist 212:1083–1093.

Conant, G. C., J. A. Birchler, and J. C. Pires, 2014. Dosage, duplication, and diploidization: clarifying the interplay of multiple models for duplicate gene evolution over time. Current Opinion in Plant Biology 19:91–98.

Defoort, J., Y. Van de Peer, and L. Carretero-Paulet, 2019. The evolution of gene duplicates in angiosperms and the impact of protein-protein interactions and the mechanism of duplication. Genome Biology and Evolution.

Deng, M., Y. Dong, Z. Zhao, Y. Li, and G. Fan, 2017. Dissecting the proteome dynamics of the salt stress induced changes in the leaf of diploid and autotetraploid paulownia fortunei. PloS one 12:e0181937.

Dong, S., V. Lau, R. Song, M. Ierullo, E. Esteban, Y. Wu, T. Sivieng, H. Nahal, A. Gaudinier, A. Pasha, et al., 2019. Proteome-wide, structure-based prediction of protein-protein interactions/new molecular interactions viewer. Plant Physiology 179:1893–1907.

Doyle, J. J. and J. E. Coate, 2019. Polyploidy, the nucleotype, and novelty: The impact of genome doubling on the biology of the cell. International Journal of Plant Sciences 180:1–52.

Edger, P. P., J. C. Hall, A. Harkess, M. Tang, J. Coombs, S. Mohammadin, M. E. Schranz, Z. Xiong, J. Leebens-Mack, B. C. Meyers, et al., 2018. Brassicales phylogeny inferred from 72 plastid genes: A reanalysis of the phylogenetic localization of two paleopolyploid events and origin of novel chemical defenses. American Journal of Botany 105:463–469.

Edger, P. P. and J. C. Pires, 2009. Gene and genome duplications: the impact of dosage-sensitivity on the fate of nuclear genes. Chromosome Research 17:699.

Fan, G., L. Wang, Y. Dong, Z. Zhao, M. Deng, S. Niu, X. Zhang, and X. Cao, 2017. Genome of paulownia (paulownia fortunei) illuminates the related transcripts, mirna and proteins for salt resistance. Scientific reports 7:1285.

Freeling, M., 2009. Bias in plant gene content following different sorts of duplication: tandem, whole-genome, segmental, or by transposition. Annual Review of Plant Biology 60:433–453.

Gabaldón, T. and E. V. Koonin, 2013. Functional and evolutionary implications of gene orthology. Nature Reviews Genetics 14:360.

Gout, J.-F. and M. Lynch, 2015. Maintenance and loss of duplicated genes by dosage subfunctionalization. Molecular biology and evolution 32:2141–2148.

Grob, S. and U. Grossniklaus, 2017. Chromosome conformation capture-based studies reveal novel features of plant nuclear architecture. Current Opinion in Plant Biology 36:149–157.

Grob, S., M. W. Schmid, and U. Grossniklaus, 2014. Hi-c analysis in arabidopsis identifies the knot, a structure with similarities to the flamenco locus of drosophila. Molecular Cell 55:678–693.

Guo, M., D. Davis, and J. A. Birchler, 1996. Dosage effects on gene expression in a maize ploidy series. Genetics 142:1349–1355.

Hakes, L., J. W. Pinney, S. C. Lovell, S. G. Oliver, and D. L. Robertson, 2007. All duplicates are not equal: the difference between small-scale and genome duplication. Genome Biology 8:R209.

Hou, J., X. Shi, C. Chen, M. S. Islam, A. F. Johnson, T. Kanno, B. Huettel, M.-R. Yen, F.-M. Hsu, T. Ji, et al., 2018. Global impacts of chromosomal imbalance on gene expression in arabidopsis and other taxa. Proceedings of the National Academy of Sciences 115:E11321–E11330.

Katju, V. and U. Bergthorsson, 2018. Old trade, new tricks: insights into the spontaneous mutation process from the partnering of classical mutation accumulation experiments with high-throughput genomic approaches. Genome Biology and Evolution 11:136–165.

Konrad, A., S. Flibotte, J. Taylor, R. H. Waterston, D. G. Moerman, U. Bergthorsson, and V. Katju, 2018. Mutational and transcriptional landscape of spontaneous gene duplications and deletions in caenorhabditis elegans. Proceedings of the National Academy of Sciences 115:7386–7391.

Langham, R. J., J. Walsh, M. Dunn, C. Ko, S. A. Goff, and M. Freeling, 2004. Genomic duplication, fractionation and the origin of regulatory novelty. Genetics 166:935–945.

Liu, C., Y.-J. Cheng, J.-W. Wang, and D. Weigel, 2017. Prominent topologically associated domains differentiate global chromatin packing in rice from arabidopsis. Nature plants 3:742.

Love, M. I., W. Huber, and S. Anders, 2014. Moderated estimation of fold change and dispersion for rna-seq data with deseq2. Genome Biology 15:550.

Lynch, M. and J. S. Conery, 2000. The evolutionary fate and consequences of duplicate genes. Science 290:1151–1155.

Lynch, M. and J. S. Conery, 2003. The evolutionary demography of duplicate genes. Pp. 35–44, in Genome Evolution. Springer.

Panchy, N., M. Lehti-Shiu, and S.-H. Shiu, 2016. Evolution of gene duplication in plants. Plant Physiology 171:2294–2316.

Papp, B., C. Pal, and L. D. Hurst, 2003. Dosage sensitivity and the evolution of gene families in yeast. Nature 424:194.

Pertea, M., D. Kim, G. M. Pertea, J. T. Leek, and S. L. Salzberg, 2016. Transcript-level expression analysis of rna-seq experiments with hisat, stringtie and ballgown. Nature Protocols 11:1650.

Pirrello, J., C. Deluche, N. Frangne, F. Gevaudant, E. Maza, A. Djari, M. Bourge, J.-P. Renaudin, S. Brown, C. Bowler, et al., 2018. Transcriptome profiling of sorted endoreduplicated nuclei from tomato fruits: how the global shift in expression ascribed to dna ploidy influences rna-seq data normalization and interpretation. The Plant Journal 93:387–398.

Qian, W., B.-Y. Liao, A. Y.-F. Chang, and J. Zhang, 2010. Maintenance of duplicate genes and their functional redundancy by reduced expression. Trends in Genetics 26:425–430.

Qiao, X., H. Yin, L. Li, R. Wang, J. Wu, J. Wu, and S. Zhang, 2018. Different modes of gene duplication show divergent evolutionary patterns and contribute differently to the expansion of gene families involved in important fruit traits in pear (pyrus bretschneideri). Frontiers in Plant Science 9:161.

Ravi, M. and S. W. Chan, 2010. Haploid plants produced by centromere-mediated genome elimination. Nature 464:615.

Riddle, N. C., A. Kato, and J. A. Birchler, 2006. Genetic variation for the response to ploidy change in zea mays l. Theoretical and Applied Genetics 114:101–111.

Robinson, D. O., J. E. Coate, A. Singh, L. Hong, M. Bush, J. J. Doyle, and A. H. Roeder, 2018. Ploidy and size at multiple scales in the arabidopsis sepal. The Plant Cell 30:2308–2329.

Rodgers-Melnick, E., S. P. Mane, P. Dharmawardhana, G. T. Slavov, O. R. Crasta, S. H. Strauss, A. M. Brunner, and S. P. DiFazio, 2012. Contrasting patterns of evolution following whole genome versus tandem duplication events in populus. Genome Research 22:95–105.

Rogers, R. L., L. Shao, and K. R. Thornton, 2017. Tandem duplications lead to novel expression patterns through exon shuffling in drosophila yakuba. PLoS Genetics 13:e1006795.

Schlapfer, P., P. Zhang, C. Wang, T. Kim, M. Banf, L. Chae, K. Dreher, A. K. Chavali, R. Nilo-Poyanco, T. Bernard, et al., 2017. Genome-wide prediction of metabolic enzymes, pathways and gene clusters in plants. Plant Physiology Pp. pp–01942.

Schnable, J. C., N. M. Springer, and M. Freeling, 2011. Differentiation of the maize subgenomes by genome dominance and both ancient and ongoing gene loss. Proceedings of the National Academy of Sciences 108:4069–4074.

Schnable, J. C., X. Wang, J. C. Pires, and M. Freeling, 2012. Escape from preferential retention following repeated whole genome duplications in plants. Frontiers in Plant Science 3:94.

Soltis, D. E., B. B. Misra, S. Shan, S. Chen, and P. S. Soltis, 2016. Polyploidy and the proteome. Biochimica et Biophysica Acta (BBA)-Proteins and Proteomics 1864:896–907.

Spoelhof, J. P., P. S. Soltis, and D. E. Soltis, 2017. Pure polyploidy: closing the gaps in autopolyploid research. Journal of Systematics and Evolution 55:340–352.

Springer, M., J. S. Weissman, and M. W. Kirschner, 2010. A general lack of compensation for gene dosage in yeast. Molecular Systems Biology 6:368.

Steige, K. A., J. Reimegård, D. Koenig, D. G. Scofield, and T. Slotte, 2015. Cis-regulatory changes associated with a recent mating system shift and floral adaptation in capsella. Molecular biology and evolution 32:2501–2514.

Stupar, R. M., P. B. Bhaskar, B. S. Yandell, W. A. Rensink, A. L. Hart, S. Ouyang, R. E. Veilleux, J. S. Busse, R. J. Erhardt, C. R. Buell, et al., 2007. Phenotypic and transcriptomic changes associated with potato autopolyploidization. Genetics 176:2055–2067.

Sun, J., Y. Shi, and E. Yildirim, 2019. The nuclear pore complex in cell type-specific chromatin structure and gene regulation. Trends in Genetics.

Taggart, J. C. and G.-W. Li, 2018. Production of protein-complex components is stoichiometric and lacks general feedback regulation in eukaryotes. Cell systems 7:580–589.

Tasdighian, S., M. Van Bel, Z. Li, Y. Van de Peer, L. Carretero-Paulet, and S. Maere, 2017. Reciprocally retained genes in the angiosperm lineage show the hallmarks of dosage balance sensitivity. The Plant Cell Pp. tpc–00313.

Visger, C. J., G. K.-S. Wong, Y. Zhang, P. S. Soltis, and D. E. Soltis, 2019. Divergent gene expression levels between diploid and autotetraploid tolmiea relative to the total transcriptome, the cell, and biomass. American Journal of Botany 106:280–291.

Wang, C., C. Liu, D. Roqueiro, D. Grimm, R. Schwab, C. Becker, C. Lanz, and D. Weigel, 2015. Genome-wide analysis of local chromatin packing in arabidopsis thaliana. Genome Research 25:246–256.

Wang, Y., X. Tan, and A. H. Paterson, 2013. Different patterns of gene structure divergence following gene duplication in arabidopsis. BMC Genomics 14:652.

Wang, Y., X. Wang, H. Tang, X. Tan, S. P. Ficklin, F. A. Feltus, and A. H. Paterson, 2011. Modes of gene duplication contribute differently to genetic novelty and redundancy, but show parallels across divergent angiosperms. PloS One 6:e28150.

Wang, Z., G. Fan, Y. Dong, X. Zhai, M. Deng, Z. Zhao, W. Liu, and Y. Cao, 2017. Implications of polyploidy events on the phenotype, microstructure, and proteome of paulownia australis. PloS one 12:e0172633.

Xie, T., F.-G. Zhang, H.-Y. Zhang, X.-T. Wang, J.-H. Hu, and X.-M. Wu, 2019. Biased gene retention during diploidization in Brassica linked to three-dimensional genome organization. Nature Plants 5:822–832.

Yan, L., G. Fan, M. Deng, Z. Zhao, Y. Dong, and Y. Li, 2017. Comparative proteomic analysis of autotetraploid and diploid paulownia tomentosa reveals proteins associated with superior photosynthetic characteristics and stress adaptability in autotetraploid paulownia. Physiology and molecular biology of plants 23:605–617.

Yao, H., A. Kato, B. Mooney, and J. A. Birchler, 2011. Phenotypic and gene expression analyses of a ploidy series of maize inbred oh43. Plant molecular biology 75:237–251.

Yu, Z., G. Haberer, M. Matthes, T. Rattei, K. F. Mayer, A. Gierl, and R. A. Torres-Ruiz, 2010. Impact of natural genetic variation on the transcriptome of autotetraploid arabidopsis thaliana. Proceedings of the National Academy of Sciences 107:17809–17814.

Zhang, J., Y. Liu, E.-H. Xia, Q.-Y. Yao, X.-D. Liu, and L.-Z. Gao, 2015. Autotetraploid rice methylome analysis reveals methylation variation of transposable elements and their effects on gene expression. Proceedings of the National Academy of Sciences 112:E7022–E7029.

Zhou, J., B. Lemos, E. B. Dopman, and D. L. Hartl, 2011. Copy-number variation: the balance between gene dosage and expression in drosophila melanogaster. Genome Biology and Evolution 3:1014–1024.

Zhu, N., P. S. Soltis, D. E. Soltis, S. Chen, and J. Koh, 2012. Proteomics and mass spectrometry of tragopogon polyploid evolution. Journal of biomolecular techniques: JBT 23:S50.

